# Dorsal and median raphe neuronal firing dynamics characterized by non-linear metrics

**DOI:** 10.1101/2023.05.23.541902

**Authors:** Claudia Pascovich Rognoni, Diego Serantes, Alejo Rodriguez, Diego Mateos, Joaqúın González, Diego Gallo, Mayda Rivas, Andrea Devera, Patricia Lagos, Nicolás Rubido, Pablo Torterolo

**Author notes:** Corresponding author: C.P. Shared Senior authors (Nicolás Rubido, Pablo Torterolo).

## Abstract

The dorsal (DRN) and median (MRN) raphe are the main serotonergic nuclei, being implicated in sleep and mood regulation. The DRN is mainly serotonergic, where neurons have regular spiking activity, slow firing rate (FR), and long action potential duration (APD). The MRN is divided in a median serotonergic region and a paramedian region, containing principally GABAergic neurons, resulting in more diverse neurochemical and electrophysiological features. In the present study, we aimed to enrich the characterization of the raphe nuclei neurons by using non-linear metrics. This was done by analyzing the neuronal basal firing profile in both nuclei of urethane-anesthetized rats using Ordinal Patterns (OP) Entropy, Bins Entropy, and Permutation Lempel-Ziv Complexity (PLZC). In a first step, we found that typical linear metrics – such as FR, coefficient of variation (CV), and APD – fail to distinguish between MRN and DRN neurons, while OP entropy is significantly different between these nuclei. We also found that the FR has a strong linear relationship with CV, Bins Entropy, and PLZC. Similarly, CV has a strong correlation with FR and Bins Entropy, whereas PLZC shows a strong linear fit with Bins Entropy. However, OP Entropy has either a weak or no linear relationship with the rest of the metrics tested, suggesting that OP Entropy is a good metric to differentiate neuronal firing profiles. In a second step, we studied how these metrics are affected by the oscillatory properties of the firing patterns. We found that all metrics are sensitive to rhythmicity – with the exception of OP Entropy. Again, this highlights OP Entropy as a powerful and useful quantity for the characterization of neuronal discharge patterns.

## 1 Introduction

The raphe nuclei are located in the brain stem along its mid-line [1]. Among these nuclei, and as part of the serotonergic system, the Dorsal Raphe Nucleus (DRN) and Medial Raphe Nucleus (MRN) are the main producers of serotonin – a widely distributed neuro-chemical in vertebrates’ central nervous system. DRN contains the greatest density of serotonergic neurons, with around 11500 in the rat, and MRN follows it with approximately 1100 [1]. A large body of evidence supports the role of the DRN and MRN in an extensive array of important functions, including stress response [2–5], pain control [6–9], reproductive functions and behaviour [10], food intake and obesity [11, 12], aggressiveness [13, 14], social interaction [15], motivation and reward [16, 17], fear [18, 19], learning and memory [20–23], motor activity [12], and the sleep-wake cycle physiology [24–26]. Additionally, the raphe nuclei play an important role in the physiopathology of several diseases, including major depression [27–29], anxiety, panic, obsessive compulsive disorder, eating disorders, phobias, drug addiction, and post-traumatic stress [30].

To date, most studies characterising neuronal groups using electrophysiological extracellular recordings focus on: the frequency of neuronal discharge, the rhythmicity (measured by the auto-correlation of events), the regularity (measured by the coefficient of variation), and the phase coherence (or phase locking) with hippocampal, cortical or other biological rhythms [31–39]. For example, serotonergic neurons from the DRN have been described as clock-like, exhibiting spontaneous activity, a slow 1 − 5 *Hz* firing-rate [40], regular spiking activity (coefficients of variation close to zero), wide action potentials (> 1.4 *ms*), and to have rhythmic patterns of discharge [40, 41], suggesting a “neuronal signature” of this neuro-chemical group. However, there are serotonergic neurons with different electro–physiological properties as well as non-serotonergic neurons in the DRN. In addition, neurons from the MRN have heterogeneous firing patterns [38, 42–45]. Therefore, there is a need to find new characteristics in the neuronal recordings in order to better differentiate between different neuronal groups, and in the case of the present paper, between DRN and MRN neurons.

The classical characterisation of neurons mainly relies on analysing the salient characteristics of the distribution of inter-spike intervals (such as the distribution’s mode and coefficient of variation) and applying linear methods (such as the auto-correlation or Z-coherence), which miss the non-linear components that are naturally present in the neuronal dynamics. Also, it is worth highlighting that some reports describe neuronal characteristics with redundant metrics; for example, mean firing-rate and mean inter-spike interval (being the inverse of each other), or the auto-correlation and the spectral components (being related by their Fourier transform), which provide no new information about the spiking characteristics.

Non-linear methods can address this problem and may reveal complementary information from the recordings – which has been the case for the electroencephalogram (EEG) [46–49]. So, we propose to use non-linear methods to extend the characterization of neuronal activity in order to improve the differentiation between functional groups of neurons. With this aim, in the present study we test these methods on the firng profile of the DRN and MRN neurons, which are known to be different in their neuronal phenotypes and localization. We used Permutation Entropy, Permutation Lempel-Ziv, and the entropy from the time-series distribution of amplitudes (i.e., Bin Entropy), and compared them with classical electrophysiological methods, which include firing frequency rate (FR), action potential duration (APD), and coefficient of variation (CV). These metrics are tested to assess whether they can distinguish the activity of DRN and MRN neurons, which are known to be different in their neuronal phenotypes and localization.

## 2 Experimental procedures

Sixty four male Wistar rats (250-310 *gr*) were used in this study. They were obtained from URBE (Reagents and Biomodels Experimentation Unit), School of Medicine, Universidad de la República. The animals were maintained with food and water available *ad libitum* and kept under controlled conditions (temperature 22 ± 2 °*C*, 12-h day–night cycle, lights on at 7:00 A.M.). All of the experimental procedures were conducted in accordance with the Guide for the Care and Use of Laboratory Animals (8th edition, National Academy Press, Washington DC, 2010) and approved by the Institutional Animal Care Commission (Exp. № 070153-000841-18). Adequate measures were taken to minimize pain, discomfort or stress of the animals, and all efforts were made to use the minimal number of animals necessary to produce reliable scientific data.

### 2.1 Recording procedures

Rats were anesthetized with urethane (1.5 *gr/kg*, i.p.) and positioned in a stereotaxic frame (David Kopf Instruments, USA). Following a scalp incision, skull landmarks were visualized and coordinates were determined from Paxinos and Watson, 2008 [50]. A small hole was drilled in the skull for unit recording of DRN and MRN neurons (AP − 8 *mm*, L 2.6 *mm*, H 6-7 for DRN and H 7-9 for MRN; from Bregma). Micropipettes were lowered at an angle of 26° for DRN and 20° for MRN to avoid the sagittal vein. Extracellular neuronal recordings were carried out using standard procedures with glass micropipette of 10-20 *M* Ω, filled with 2M NaCl [51–54] or with 2% Neurobiotin (Nb, Vector Laboratories) in 0.5M NaCl solution [55]. Two screw electrodes were placed in the frontal and parietal cortices, and two twisted nichrome electrodes were placed in the dorsal hippocampus (coordinates: AP, -4.4 *mm*; L, -2.6 *mm*; H, 3.4 *mm*) to monitor the electrocorticogram (ECoG).

Neuronal signals were amplified by an AC-coupled amplifier (Dagan 2400A), filtered between 0.3 *Hz*-10 *kHz* and digitized at 20 *kHz*. Single unit activity was acquired and processed using Spike 2 software (Cambridge Electronic Design, UK). The baseline discharge of raphe neurons was recorded for 3 to 10 min. Neurobiotin was administered by iontophoresis with the following protocol: anodic current, 5 *nA*, 200 *ms* on / 200 *ms* off, for 5 minutes [53]. In order to visualize the ECoG, the signal was amplified (×1000), filtered (0.1–100 *Hz*), acquired (512 *Hz*, 16 bits) and processed with Spike 2. The ECoG was used to check the electrocortical state of anesthesia. Only the neurons recorded in the slow wave state or Non-REM urethane [56] were selected for the analyses.

### 2.2 Histological and immunofluorescence procedures

Two types of procedures were performed depending on the experiment. First, in the experiments that the rats did not receive Nb, after finishing the recording experiments and dissecting the brain, it was left immersed in 10% formalin for 48 hours and then was sectioned into slices of 100 *μm* using a vibratome in order to determine the location of the recording electrodes. The path of the micropipette and the location were recognized by light microscopy, and photographs were taken with a histological magnifying glass.

Second, at the end of those experiments in which a neuron was labeled with Nb, the animals were perfused transcardially with NaCl 0.9% heparinized and then 4% paraformaldehyde (PFA). The brains were removed and post-fixation was performed in 4% PFA for 24 hours. Afterwards they were left in immersion in 30% sucrose for 48 hours. They were finally cut into blocks and frozen on dry ice. Coronal sections 30 *μm* of thickness were obtained by a cryostat and stored with cryoprotection solution at −20°*C*.

The identification of the neuron labeled with Nb was performed by two immunohistochemical procedures. In the first procedure, after washing the sections with phosphate buffered saline (PBS), neurons were incubated with PBS plus 0.3% Triton X-100 (PBST) for 90 min. Then, they were incubated with 1% H2O2 for 30 min. After washing with PBS, the sections were incubated with peroxidase-avidin-biotin complex (ABC 1: 200, Vector Labs) for 120 min and then were exposed to diaminobenzidine (DAB, 0.02%) for 10 min. Then, the sections were washed again and were mounted and coated with glycerol. Finally, photomicrographs were taken in light microscope (Olympus model) to visualize the Nb labeled neuron. In the second procedure, an immunofluorescence was performed. The sections were incubated in 0.5% sodium borohydride for 25 min; thereafter, they were incubated in PBS. Then, they were incubated with streptavidin-Alexa Fluor 555 conjugate (1:5000, Molecular Probes) in PBST 0.3% for 2.5 h. Finally sections were mounted and observed under an epifluorescence microscope in order to find the neuron labeled with Nb (examples are shown in Fig. 1). In occasions, more than one neuron was labeled with Nb, but the location of the recorded neuron was clearly identified.

**Figure 1:**
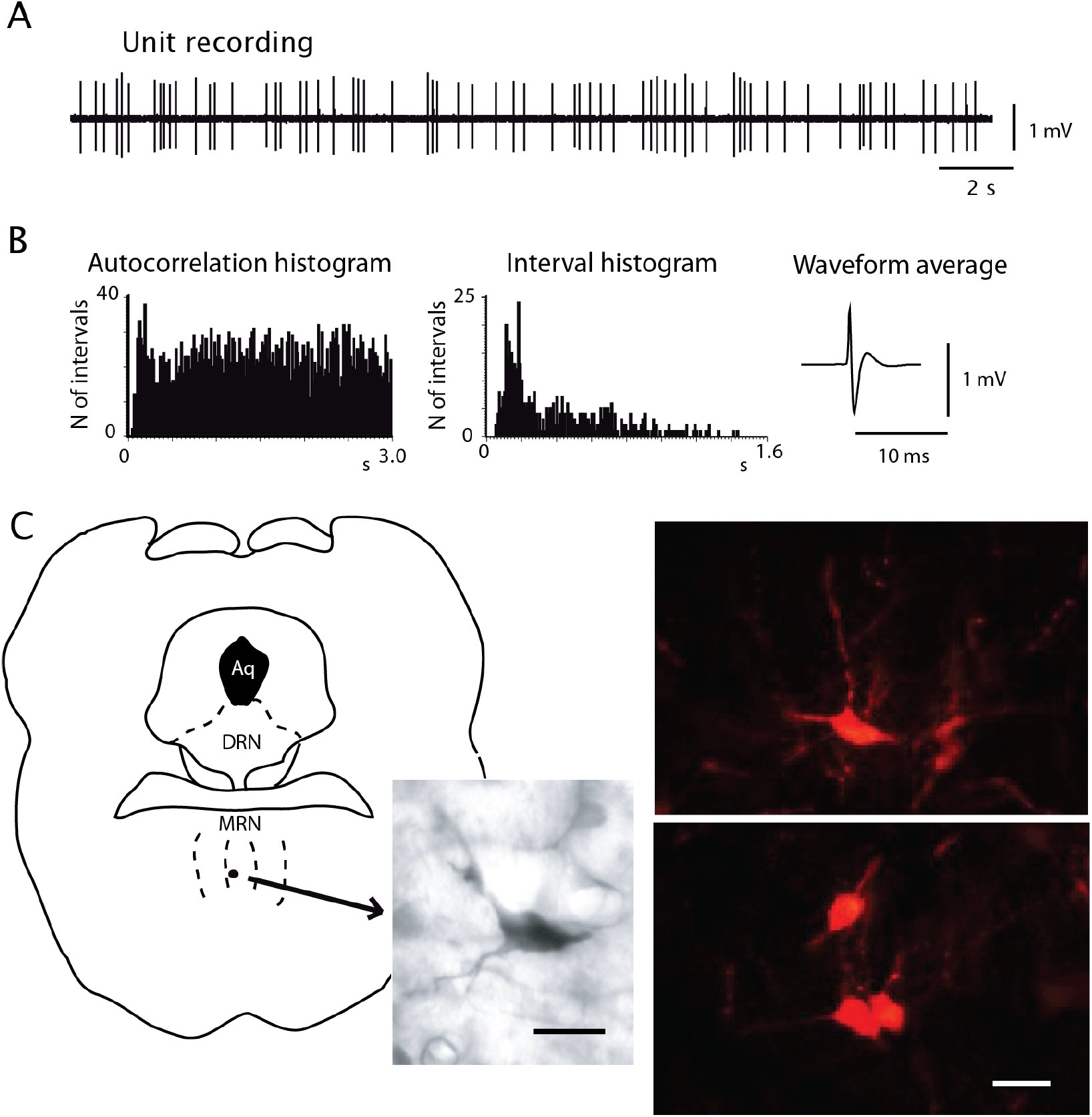
Examples of neuronal recording and identification. A) Neuronal raw recording of a MNR neuron. B) Auto-correlation histogram, interval histogram and waveform of the same neuron. C) Examples of neurons identified by neurobiotin. Antero-posterior coordinate from Bregma −8 mm according to [50]. Calibration bars: 20 *μ*m.

### 2.3 Data analysis

We sorted the single units according to amplitude and waveform criteria. We examined the lack of spikes during the refractory period (< 2 *ms*), confirming the absence of contamination by other units.

#### Classic metrics

The action potential (AP) of each neuron were averaged and analyzed in shape and duration. The APs were mostly triphasic, and the duration of the first two phases was considered as the AP duration (APD) [31]. The frequency of spontaneous activity (FR) and standard deviation was also calculated. Additionally, the pattern of spontaneous discharge with interval (IH) and autocorrelation (ACH) histograms were analyzed. The coefficient of variation (CV) was used to determine the regularity of the discharge frequency. These analyses were performed in windows of 60-300 *s* of stable activity. Neurons with burst activity were identified by looking at the raw recordings and IH.

#### Ordinal Pattern Entropy

The degree of randomness in sets of consecutive inter-spike intervals (ISI) was quantified by the Permutation Entropy [57], which we name OP entropy and is the Shannon Entropy [58] of the Ordinal Pattern (OP) encoding of the ISI sequence. Specifically, the OP Entropy is found from 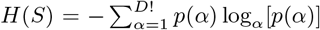, where *p*(*α*) is the probability of having the OP *α* in the encoded ISI sequence (i.e., the relative frequency of appearance of *α*) and there can be *D*! different OPs. Specifically, OPs are obtained by splitting the ISI sequence into non-overlapping sets with *D* consecutive data-points and calculating the number of permutations needed to organize the ISI magnitudes within each set in increasing order. For example, setting *D* = 2, the ISI sequence is divided into the following sets: *{x*(*t*_0_), *x*(*t*_1_)*}, {x*(*t*_1_), *x*(*t*_2_)*}*, and so on, where *x*(*t*_*i*_) is the ISI value at time *t*_*i*_. Each set is transformed into an OP by assessing whether *x*(*t*_*i*_) < *x*(*t*_*i*+1_) or *x*(*t*_*i*_) > *x*(*t*_*i*+1_), which correspond to making 0 or 1 permutation, respectively, and having *D*! = 2! different OPs. In this work we use *D* = 3, so that ISI sequences are encoded into 6 possible OPs and we have sufficient statistical power to find the probabilities *p*(*α*). Also, in order to remove possible cases where *x*(*t*_*i*_) = *x*(*t*_*i*+1_), a negligible white-noise signal is added to the ISI sequence before performing the OP encoding.

#### Bins Entropy

Is the Shannon Entropy [58] of the ISI histogram, which quantifies the degree of randomness in the magnitudes of the ISIs. Its value is obtained from 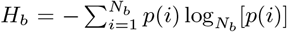, where *N*_*b*_ is the number of bins used to construct the histogram. In this work we set to *N*_*b*_ = 18 to maintain the same statistical power as with the OP entropy.

#### Permutation Lempel-Ziv complexity

Lempel-Ziv complexity (LZC) is an information measure based on the Kolmogorov complexity – the minimal “information” contained in the sequence [59]. This complexity has been used in the analysis of different types of neurophysiological signals, among others for the study of the effects of anaesthesia, seizures, and consciousness [60–62]. To estimate the complexity of a time series *𝒳* (*t*) *≡ {x*_*t*_; *t* = 1, …, *T }* we used the Lempel and Ziv scheme proposed in 1976 [63]. In this approach, a sequence *𝒳* (*t*) is parsed into a number *𝒲* of words by considering any sub-sequence that has not yet been encountered as a new word. The Lempel-Ziv complexity *c*_*LZ*_ is the minimum number of words *𝒲* required to reconstruct the information contained in the original time series. For example, the sequence 100110111001010001011 can be parsed in 7 words: 1 *·* 0 *·* 01 *·* 101 *·* 1100 *·* 1010 *·* 001011, giving a complexity *c*_*LZ*_ = 7. A way to apply the Lempel-Ziv algorithm can be found in [64]. The LZC can be normalized based in the length *T* of the discrete sequence and the alphabet length (*α*) as *𝒞*_*LZ*_ = *c*_*LZ*_ [log_*α*_ *T*]*/T*. Although, initially Lempel and Ziv developed the method for binary sequences, it could be used for any alphabet with finite length. In particular LZC could be applied over the OP discretization, which is known as *Permutation Lempel-Ziv complexity* [65].

### 2.4 Statistics

In this work, group comparisons were performed using Multilevel Bayesian models, and the null hypothesis was rejected at *p* < 0.05. We chose these models due to the hierarchical nature of the data and bimodal or multimodal and skewed nature of the distributions. A variable number of neurons (from 1 to 7) were recorded per rat, with a total of 169 neurons in 64 rats. The rat was included in the model as a random effect, and the metric to be explored (classic and non-linear metrics) as the fixed effect. All models were estimated using the open-source packages MCMCglmm v.2.30 on R v.3.6.1. ggforce-package was used for the clusters analysis.

## 3 Results

We recorded a total of 169 neurons from both nuclei, corresponding 77 to the DRN and 92 to MRN of the rat. We confirmed that the neurons were located within the limits of the corresponding nuclei by reconstruction of the micropipette tracts, or by identification of the recorded neuron with neurobiotin (Nb). Fig. 1 shows an examples of the recording, processing and recognition of Nb-labeled neurons.

The entire population of recorded neurons within the DRN (*n* = 77) display the following electrophysiological characteristics. An APD of 3.07 ± 0.37 *ms*, FR 8.45 ± 1.94 *Hz*, and CV 0.65 ± 0.079. 79% percent of these neurons exhibit uni-modal IH, 26% have a rhythmic pattern of discharge in the ACH, and 20% have a predominant interval in the ACH. Moreover, a burst firing pattern is observed in 4 neurons (5%), i.e., showing doublets or triplets with < 20 *ms* intervals and a prominent decrease in the amplitude of higher order spikes [31]. On the other hand, the recorded neurons in the MRN (*n* = 92) display the following electrophysiological characteristics. An APD of 2.56 ± 0.27 *ms*, FR 10.52 ± 1.34 *Hz*, and CV 0.76 ± 0.055. 75% of these neurons exhibit uni-modal IH, 21% a rhythmic pattern, and 35% a predominant interval in the ACH. Burst firing pattern is observed in 20 neurons (21%).

### 3.1 OP Permutation Entropy distinguishes DRN and MRN neuronal activity

We first studied whether the non-linear metrics – OP Entropy, PLZC, and Bins Entropy – can distinguish MRN from DRN neuronal dynamics compared to the classic metrics – APD, CV, and FR. For the classic metrics, it can be seen from Fig. 2 that only the CV shows a slightly non-significant trend in discriminating both nuclei (0.77 ± 0.05 for the MRN Vs. 0.61 ± 0.08 for the DRN, *p* = 0.06), whereas the FR and the APD are not significantly different between groups (10.74 ± 1.39 vs 8.72 ± 2.08, *p* = 0.30; 2.47 ± 0.29 vs 2.72 ± 0.40 *ms, p* = 0.33). For the non-linear metrics, Bins Entropy and PLZC show non-significant differences between the values from both groups (0.52 ± 0.02 Vs. 0.54 ± 0.03, *p* = 0.63, and 0.76 ± 0.005 Vs. 0.77 ± 0.008, *p* = 0.28, respectively), but the OP Entropy is significantly lower in the DRN compared to the MRN (0.9970 ± 5.714 *×* 10^*−*4^ Vs. 0.9954 ± 8.460 *×* 10^*−*4^, *p* < 0.05). This significant difference can be seen in the violin plots from Fig. 2A, which could be attributed to a sub-population of neurons with an OP Entropy lower than 0.985.

**Figure 2:**
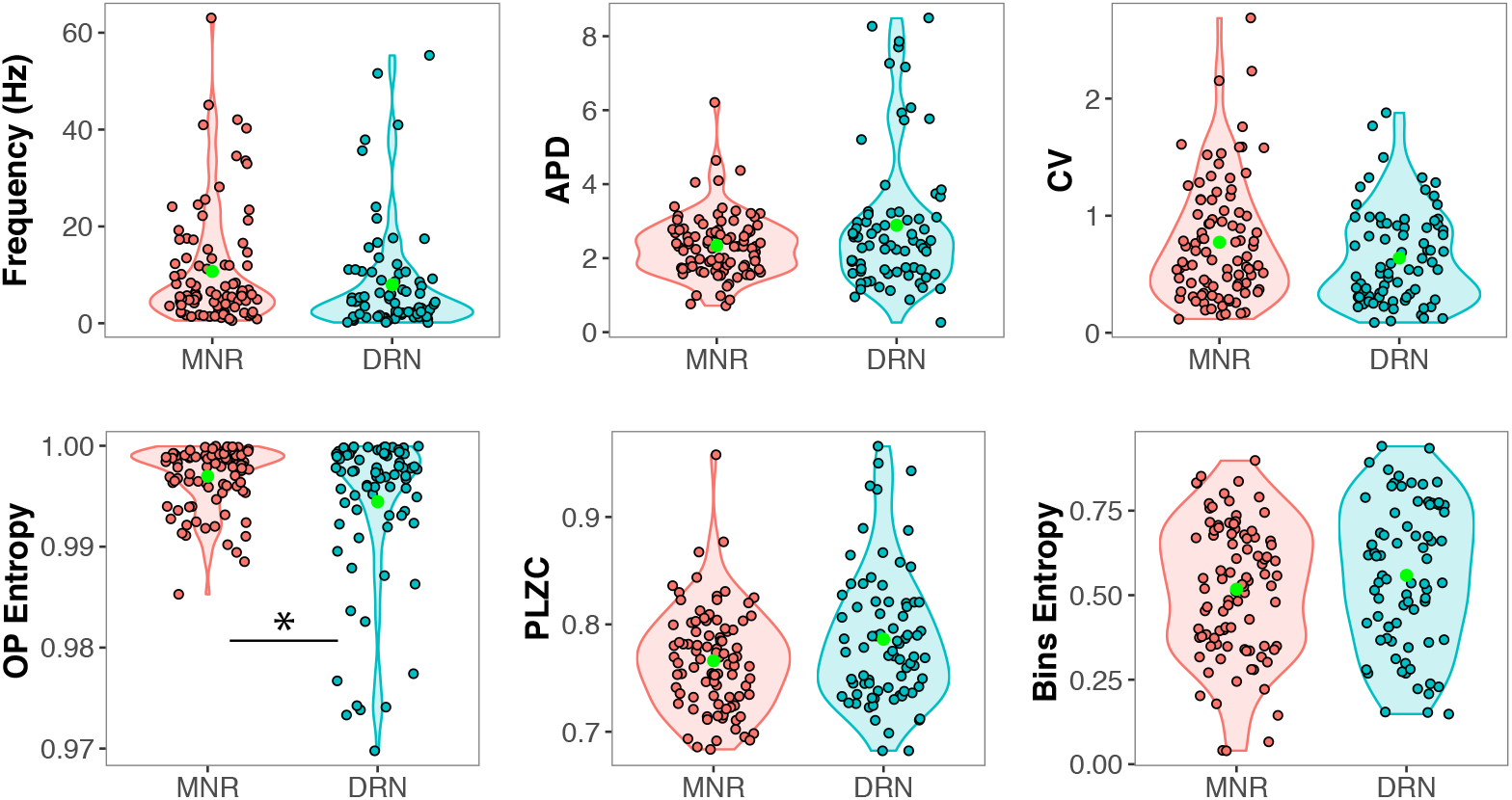
Comparison between classic metrics (top) and non-linear metrics (bottom) in distinguishing MRN and DRN neuronal sub-populations. Classic metrics cannot distinguish the nuclei, and only OP Entropy among the non-linear metrics is significantly different between both neuronal groups. OP Entropy, Ordinal Patterns Entropy; PLZC, Permutation Lempel-Ziv complexity; APD, action potential duration; CV, coefficient of variation. \**p* < .05

We also explored different paired combinations of the variables and analyzed the resultant scatter plots. For each pair, two contours are plotted surrounding the data-points which belong to the DRN (light blue) and MRN (orange); by this approach we can determine if the DRN and MRN neuronal populations are superimposed or could be separated by a specific combination of variables (Fig. 3).

**Figure 3:**
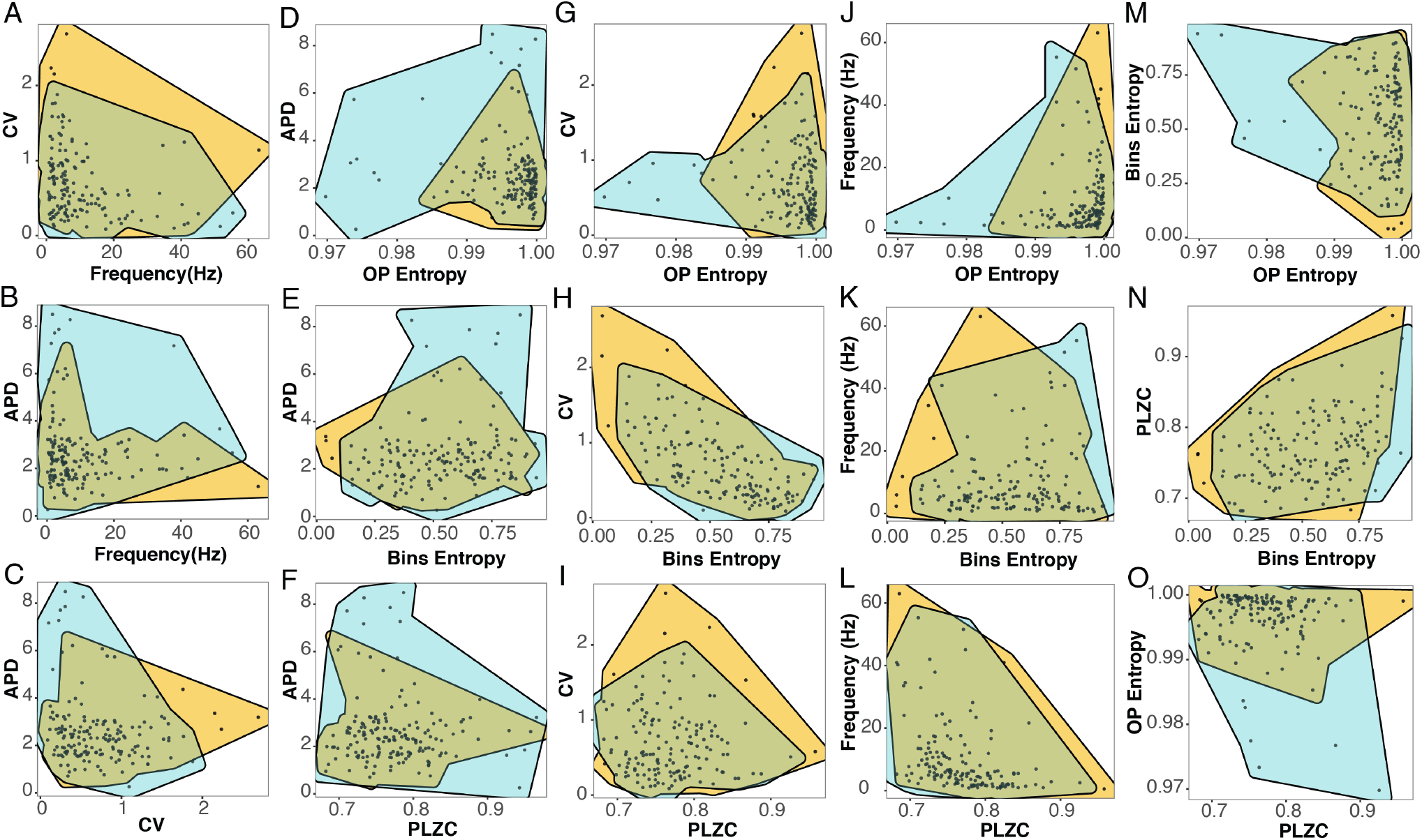
Scatter plots and cluster identification of DRN (light blue) and MRN (orange). Superimposed area is represented in brown. OP Entropy, Ordinal Patterns Entropy; PLZC, Permutation Lempel-Ziv complexity; APD, action potential duration; CV, coefficient of variation.

The combination of FR and CV does not clearly separate the two nuclei (Fig. 3A). The combination of APD with FR or CV only partially separates both nuclei, where the DRN shows a group with high APD and low FR (Figs. 3B), and the MRN a group with low APD and high CV(3C).

When comparing the APD with non-linear metrics (Figs. 3D, 3E, and 3F), the combinations can separate some DRN neurons (represented in the figure in light blue) out from the superimposed brown area. For example, it can be seen (Fig. 3D) that neurons with OP Entropy lower than 0.985 have a short APD and that these neurons belong to the DRN. By contrast, MRN neurons show OP Entropy higher than 0.985. For Bins Entropy and PLZC the non-superimposed area predominates in the upper part of the plots, for the neurons with larger APD from the DRN (Figs. 3E, and 3F).

When looking at the relationship between the CV and non-linear metrics (Figs. 3G, 3H, and 3I), results vary. The combination of CV and OP entropy can distinguish some DRN from MRN neurons (Figs. 3G). There is a group of neurons with OP Entropy lower than 0.985 that purely contain DRN neurons with small CV, whereas for higher values of OP Entropy both neuronal population are intermingled (as in Fig. 2G).

On the other hand, there are no noticeable differences between DRN and MRN neurons when looking at the combination of CV and Bins Entropy or PLZC (Figs. 3H and 3I). Similarly, the combination of FR with non-linear metrics (Figs. 3J, 3K, and 3L) shows partial separation between both nuclei when combined with OP Entropy (Fig. 3J), where entropy values below 0.98 contain solely neurons from the DRN with low FR.

Finally, the combinations between the entropies (Figs. 3M, 3N, and 3O) can separate some of the neurons from the DRN and MRN populations. When using OP entropy and Bins entropy (Fig. 3M), a light blue area can be seen containing neurons with low OP Entropy and high Bins Entropy; and when using PLZC with OP Entropy DRN neurons with low OP Entropy, shown in light blue, can be visualized but display varied PLZC values (Fig. 3O).

From these results, we can conclude that the APD combined with non-linear metrics, shown in Figs. 3D, 3E, and 3F, are better in separating the two raphe neuronal populations. Additionally, the OP Entropy can also separate the discharge pattern of these nuclei populations in combination with other variables including CV, FR, Bins Entropy and PLZC.

### 3.2 Relationship between classic and non-linear metrics

In order to explore in more detail the relationship between the different metrics for both nuclei, we plotted all possible paired combinations of the variables and studied the relationship between them (Fig. 4 and Table 1), by the use of linear and quadratic model fitting.

**Figure 4:**
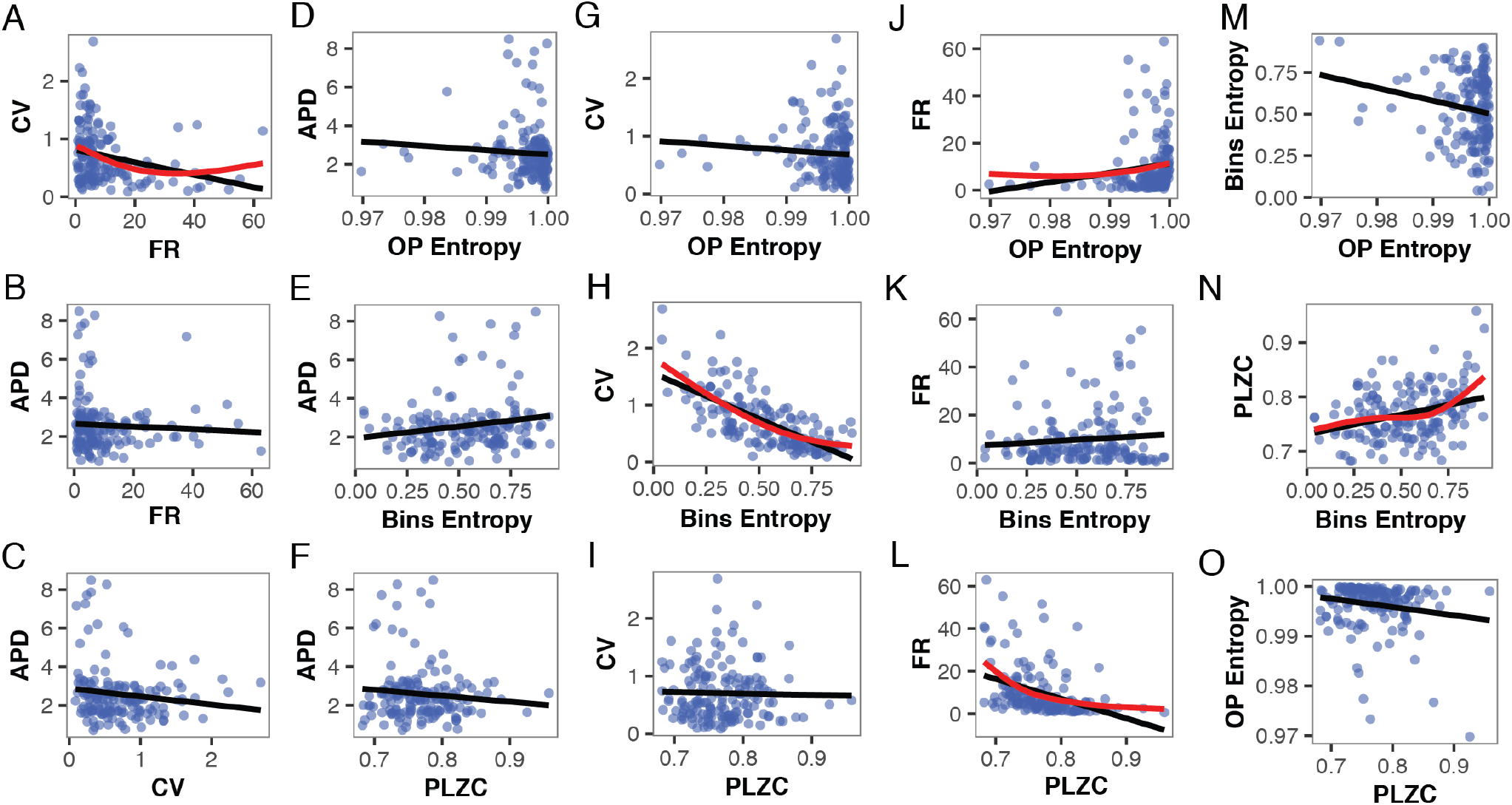
Linear relationships between all combination of variables are represented with a black line. The relationships were non-linear for some variable pairs (represented in red).

**Table 1:** Statistical comparison between metrics using linear multilevel models. \**p* < .05, \*\**p* < .005, \*\*\**p* < .001.

|  | Mean Freq. | CV | APD | OP Entropy | Bins Entropy | PLZC |
| --- | --- | --- | --- | --- | --- | --- |
| Mean Freq. |  | < 0.001*** | 0.616 | 0.044* | 0.464 | < 0.001 * ** |
| CV | < 0.001*** |  | 0.14 | 0.328 | < 0.001*** | 0.756 |
| APD | 0.616 | 0.14 |  | 0.346 | 0.05 | 0.198 |
| OP Entropy | 0.044* | 0.328 | 0.346 |  | 0.024* | 0.022* |
| Bins Entropy | 0.464 | < 0.001*** | 0.05. | 0.024* |  | < 0.001*** |
| PLZC | < 0.001*** | 0.756 | 0.198 | 0.022* | < 0.001*** |  |

For the classic metrics, there is a statistically significant negative linear relationship between the FR and the CV (*p* < 0.001, Fig. 4A), but there is neither a linear relationship between the FR and APD, nor between the CV and the APD (*p* = 0.61 in Fig. 4B and *p* = 0.14 in Fig. 4C).

When looking at a linear relationship between the APD and the non-linear metrics, it can be seen that none of them show a significant fit (Figs. 4D, 4E, and 4F). In the case of the CV, there is a significant negative linear relationship with the Bins entropy (*p* < 0.001, Fig. 4H) but no significant linear relationship with OP Entropy or PLZC (Figs. 4G and 4I). The mean FR shows a negative linear relationship with the OP Entropy and with PLZC (*p* = 0.044 and *p* < 0.001 in Figs. 4J and 4L, respectively). However, the FR and the Bins Entropy are not linearly related. One important thing to note is that the relationship between FR and OP Entropy is actually non-linear, since all the neurons with OP Entropy lower than 0.99 show a low FR (lower than 10 *Hz*). However, in the group with higher OP entropy, a wide range of FR (from 0.55 to 53 *Hz*) can be found (Fig. 4J).

Regarding the correlation between the non-linear metrics, it can be seen that Bins Entropy and OP Entropy show a weak negative linear relationship (*p* = 0.024, Fig. 4M) as well as OP Entropy and PLZC (*p* = 0.022, Fig. 4O). By contrast, PLZC and Bins Entropy are strongly positively related (*p* < 0.001, Fig. 4N).

In Table 1 we summarize the statistical results for all the linear combinations of the classic and non-linear metrics. Then, for the significant pairs in the linear models, we performed model selection using Bayes Factors (BF) to address whether the relationship between the pair of variables is actually explained better by a quadratic function. We found that for all model comparison showed the supremacy of quadratic fit (Table 2).

**Table 2:** Selection between linear and non-linear models among different combination of metrics. Mixed effect models were applied to pair of metrics, including the rat as a random effect. Bayes Factors (BF) were used to decide between quadratic and linear fits. With the exception of PLZC/Bin Entropy, all model comparisons showed the supremacy of the quadratic fit, where asterisks indicate their substantial evidence (BF > 5).

| Metrics pairs | Model | BF |
| --- | --- | --- |
| FR / CV | linear |  |
| | quadratic * | $1.32 \times 10^{220}$ |
| OP Entropy / FR | linear |  |
|  | quadratic * | 3932.6 |
| FR / PLZC | linear |  |
|  | quadratic * | 987.83 |
| CV / Bin Entropy | linear |  |
|  | quadratic * | 33.72 |
| Bin Entropy / OP Entropy | linear |  |
|  | quadratic * | 80.98 |
| OP Entropy / PLZC | linear |  |
|  | quadratic * | 65 |
| PLZC / Bin Entropy | linear |  |
|  | quadratic | 0.22 |

### 3.3 Sensitivity of the classic and non-linear metrics to rhythmic pattern

In order to describe in more detail the population of raphe nuclei comparing the classic and non-linear metrics, we studied how these metrics are influenced by the rhythmicity of the neuronal activity. Specifically, some neurons in the DRN and MRN fire predominantly in specific frequencies (1 up to 60 *Hz*, Fig. 5A), repeat their firing patterns periodically and have an ACH with regular peaks (Fig. 5B). It is interesting to note that, rhythmic neurons of the MRN discharge predominantly in the theta rhythm and the proportion of neurons firing within this frequency range is lower in the DRN (Fig. 5A). This results are in line with previous evidence which shows that a subgroup of neurons, discharged rhythmically in synchrony with the hippocampal theta rhythm; the synchronization of the firing with the hippocampal theta rhythm is greater in the MRN than in the DRN [36].

**Figure 5:**
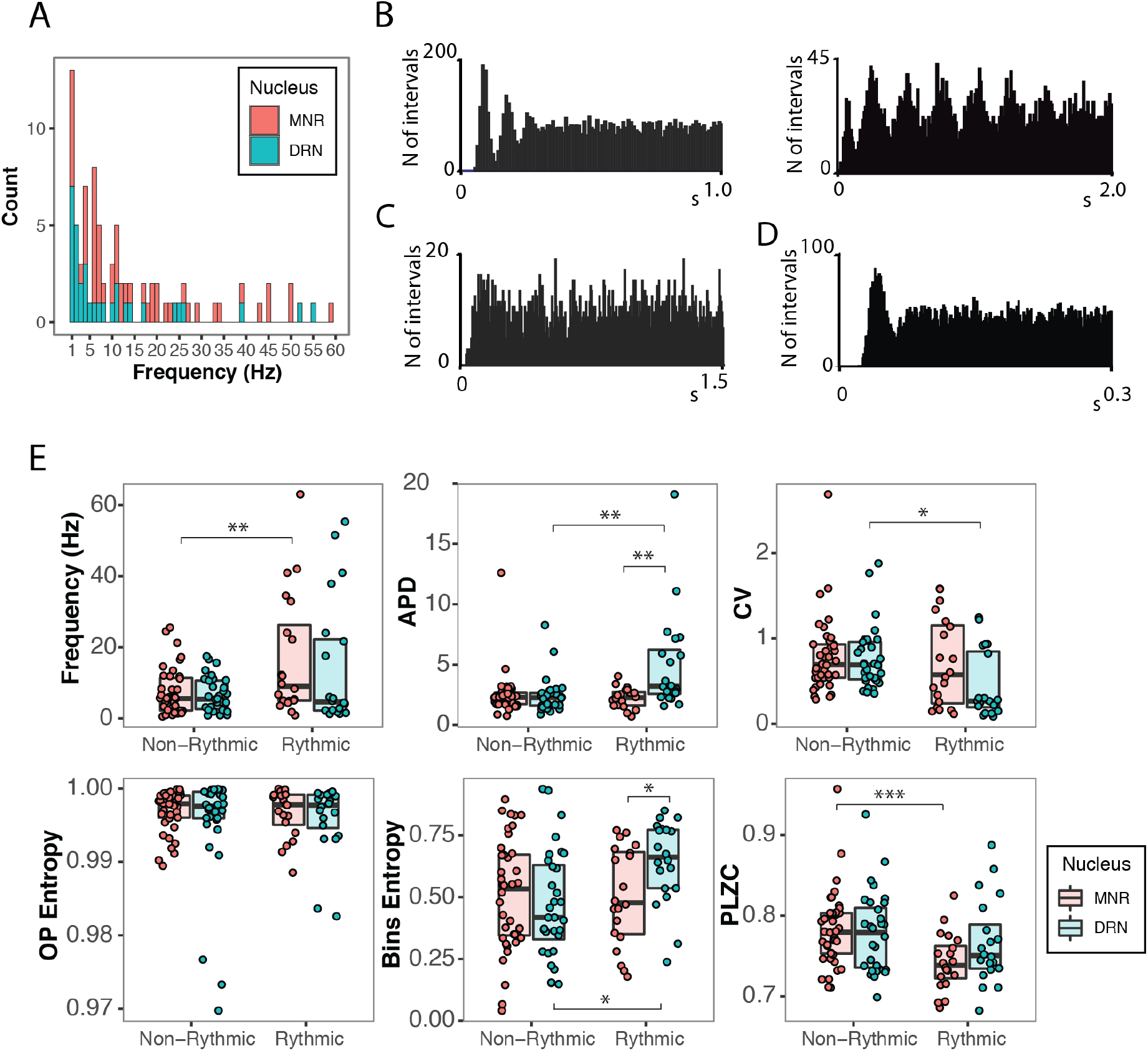
Sensitivity of the classic and non-linear metrics to rhythmic patterns in the neuronal discharge. A) Distribution of the frequencies of the rhythmic neurons of the DRN and MRN. B) ACH of rhythmic neurons. C) ACH of a non-rhythmic neuron. D) Example of a single predominant interval in the ACH. E) Relationship between classic and non-linear metrics with rhythmicity, where OP Entropy is the only metric insensitive to the rhythmic pattern. The median is represented with a horizontal black lines. *p < .05, **p < .005, ***p < .001.

Here we classified the neurons from each nucleus into two categories: rhythmic and non-rhythmic. Neurons having an ACH without defined peaks or showing only a unique predominant interval in the ACH were excluded from this analysis, as in the example of Fig. 5D. Therefore, we only compare clearly rhythmic (Fig. 5B), showing more than one peak, or clearly non-rhythmic pattern (Fig. 5C). Then, we tested if the different metrics are affected by rhythmic properties and explored the ability of each of the metrics in separating DRN and MRN within rhythmic and non-rhythmic subgroups.

When looking at the classic metrics, it can be seen from Fig. 5E that the FR, APD, and CV are modulated by rhythmicity depending on whether the neurons are from the MRN or DRN. Rhythmic neurons from the MRN group have a significantly higher FR than non-rhythmic ones (16.96±3.07 *Hz* and 7.43±1.77 *Hz* respectively, *p* = 0.008), but rhythmic and non-rhythmic neurons from the DRN have similar FR values (12.60±3.49 *Hz* and 8.07±2.11 *Hz* respectively, *p* = 0.23). On the other hand, APD is significantly larger in rhythmic neurons from the DRN than non-rhythmic ones (3.89 ± 0.50 *ms* versus 2.4661 ± 0.29 *ms*, respectively, *p* = 0.005), but the APD shows no significant differences in the rhythmicity of MRN neurons (2.23 ± 0.12 *ms* vs 2.23 ± 0.21 *ms, p* = 0.998). Similarly, rhythmic neurons from the DRN have significantly smaller CV than non-rhythmic ones (0.46 ± 0.11 and 0.74 ± 0.06 respectively, *p* = 0.01), but are undifferentiated in the MRN (0.71 ± 0.12 versus 0.78 ± 0.073, *p* = 0.54).

Regarding the non-linear metrics, it can be observed from Fig. 5E that OP Entropy is the only one insensitive to rhythmicity. For the DRN, the OP Entropy for non-rhythmic and rhythmic neurons is 0.99 ± 0.0011 and 0.99 ± 0.0019 (*p* = 0.77), respectively. For the MRN, the OP Entropy values are 0.99 ± 0.00048 and 0.99 ± 0.00082, respectively.

On the other hand, Bins Entropy significantly increases to 0.63 ± 0.059 in the rhythmic neurons of the DRN versus the non-rhytmic neurons, which hold 0.47 ± 0.035 (*p* = 0.006). However, no differences are found for the MRN, with Bins Entropies of 0.50±0.034 and 0.50±0.059 (*p* = 0.956) for non-rhythmic and rhythmic neurons, respectively. Similarly, PLZC is sensitive to rhythmic properties of the neuronal discharge, where rhythmic neurons have lower PLZC of 0.74 ± 0.011 than non-rhythmic neurons, with 0.78 ± 0.0068. However, it was only statistically significant for the MRN (*p* < 0.001). No differences were found for the DRN (0.77 ± 0.0088 Vs. 0.765 ± 0.012, *p* = 0.64).

When looking at the ability of linear and non-linear measures to differentiate between DRN and MRN neurons within rhythmic or non-rhythmic subgroups, it can be seen that only APD and Bins Entropy achieve a significant difference in their values for rhythmic neurons. APD is larger for the DRN (0.66 ± 0.058 *ms*) compared to the MRN (0.50 ± 0.039 *ms, p* = 0.012), and Bins Entropy is higher for the DRN (0.50 ± 0.041 Vs. 0.63 ± 0.059, *p* = 0.04) compared to the MRN. However, no metrics separates the nuclei when looking at the non-rhythmic subgroup of neurons.

### 3.4 Non-linear metrics are insensitive to burst patterns

Some neurons in DRN (5%) and MRN (21%) display a burst firing pattern. An example of this firing pattern is shown in Fig. 6A. In order to study how this pattern of activity can affect the classic and non-linear metrics we compared neurons with and without burst pattern activity for each of the variables and for both nuclei (Fig. 6B). We encountered that of the classic metrics only the CV is sensitive to burst activity having the neurons with burst a higher CV for DRN (1.07 ± 0.19 Vs. 0.58 ± 0.05; *p* < 0.05) as well as for the MNR (1.07 ± 5.31 Vs. 0.69 ± 1.90; *p* < 0.01). None of the other metrics are affected by the presence of burst pattern of activity (FR for DRN: 4.59 ± 5.31 Vs. 10.13 ± 1.90, *p* = 0.36; FR for MRN: 15.61 ± 5.31 Vs. 9.81 ± 1.90, *p* = 0.98; APD for DRN: 2.13 ± 1.26 Vs. 3.41 ± 0.50, p = 0.37; APD for MRN: 2.44 ± 0.34 Vs. 2.45 ± 0.16, p = 0.98; OP Entropy for DRN: 0.992 ± 3.3 *×* 10^*−*3^ Vs. 0.995 ± 0.82 *×* 10^*−*3^, p= 0.27; OP Entropy for MRN: 0.996 ± 0.77 *×* 10^*−*3^ Vs. 0.997 ± 0.35 *×* 10^*−*3^, p= 0.27; PLZC for DRN: 0.76 ± 0.25 *×* 10^*−*1^ Vs. 0.77 ± 0.63 *×* 10^*−*2^, p = 0.67; PLZC for DRN: 0.74 ± 1.17 *×* 10^*−*2^ Vs. 0.77 ± 0.54 *×* 10^*−*2^, p = 0.05; Bins Entropy for DRN: 0.9921 ± 0.11427 Vs. 0.54697 ± 0.02813, p = 0.52). Bins Entropy for MRN: 49.86 *×* 10^*−*2^ ± 0.50 *×* 10^*−*1^ Vs. 52.31 *×* 10^*−*2^ ± 0.23 *×* 10^*−*1^, p = 0.61).

**Figure 6:**
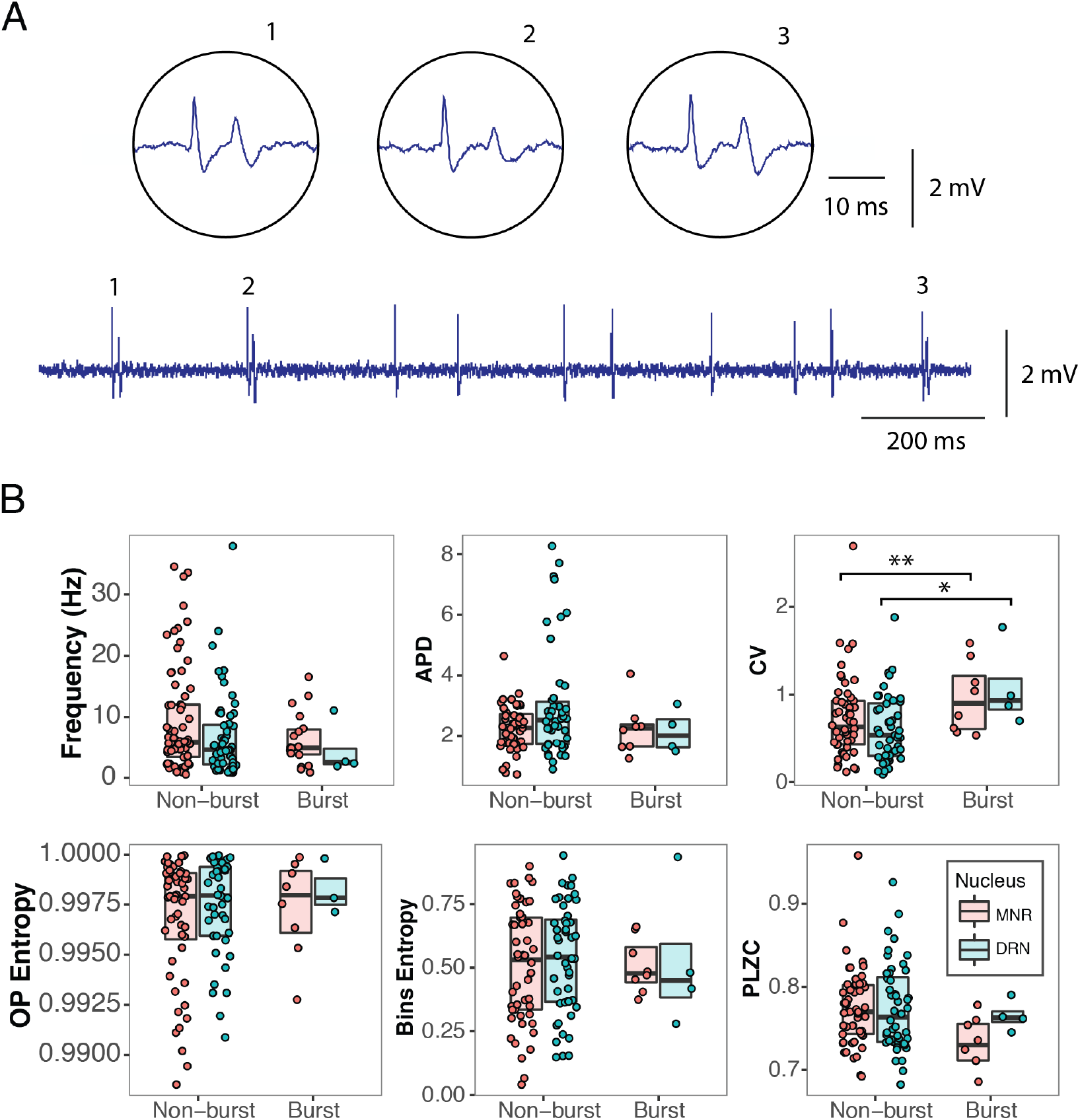
Sensitivity of the classic and non-linear metrics to the presence of bursts in the neuronal discharge. A) Example of a raw neuronal recording with bursts (indicated with numbers). A zoomed view of these bursts is shown in the circular boxes. B) Comparison of the six metrics studied on their sensitivity to bursts – non-linear metrics appear insensitive to bursts. The CV is the only metric that shows a higher value for the bursting neurons in both nuclei. The median is represented by horizontal black lines. *p < .05, **p < .005, ***p < .001.

### 3.5 A high Bins Entropy characterize the putative serotonergic neurons

Putative serotonergic neurons had been determined to have APD larger than 1.4, CV lower than 0.30, rhythmic pattern in the ACH, and low FR (< 4*Hz*) [33, 35, 66, 67]. Neurons considered as putative serotonergic were separated from the non-serotonergic, and were compared with non-linear metrics. Neurons with those characteristics were only 7 out of 66 and were only present in the DRN.

As shown in Fig. 7, putative serotonergic neurons showed a higher Bins Entropy compared with the non-serotonergic group (0.51 ± 0.03 vs 0.78 ± 0.07, *p* = 0.002). The OP Entropy and PLZC were not statistically different between those groups (OP Entropy: 0.995 ± 8.456*e −* 04 vs 0.997 ± 2.596*e −* 03, *p* = 0.42; PLZC: 0.77 ± 0.0064 vs 0.79 ± 0.019, *p* = 0.38).

**Figure 7:**
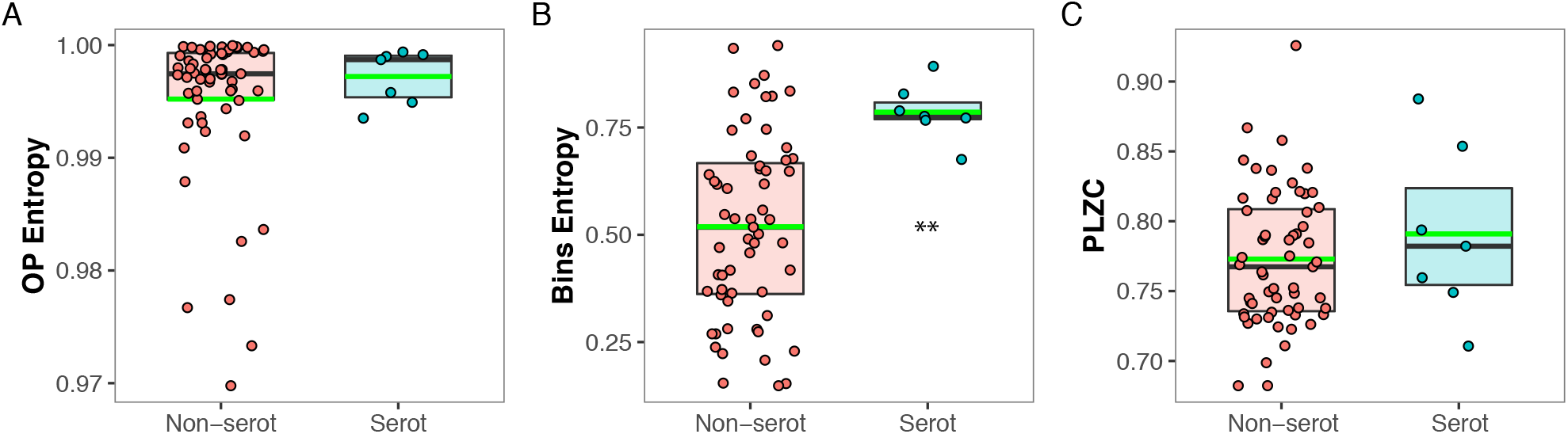
Characterisation of the putative serotonergic and non-serotonergic neurons using non-linear metrics: (A) OP Entropy, (B) Bins Entropy and (C) PLZC. Putative serotonergic neurons show high Bins Entropy whereas non-serotonergic sub-group shows a widespread distribution. OP, Ordinal Patterns; PLZC, Permutation Lempel-Ziv complexity. The median is represented by horizontal black lines and the mean with green lines. **p < .005

## 4 Discussion

In the present work, we characterize the dynamics of neuronal activity comparing DRN and MRN using classic metrics – FR, APD and CV – and the more recently developed non-linear metrics – OP Entropy, Bins Entropy and PLZC. We first summarize the evidence of the classical metrics of the DRN and MRN, and then we discuss the contribution of the non-linear metrics to the characterization of raphe neuronal dynamics.

The importance of the study of raphe neurons, specially their serotonergic neurons, lies in its implications in physiology, but mainly, in the physiopathology of mood disorders; serotoninergic neurons are the main target of currently used antidepressant drugs. Since the 1970s, several research groups have been trying to identify electrophysiological characteristics that could distinguish serotonergic neurons [31–33, 40, 66].

The DRN and MRN nuclei are by far the main source of serotonin in the brain, but the neurochemical characteristics of DRN and MRN differ significantly, having the DRN the great majority of such serotonergic neurons [1]. It is also known that GABAergic neurons are present in the DRN and MRN [68]. However, while serotonergic neurons are located mainly in the brain’s midline, GABAergic neurons are not only in the midline but also in the lateral regions of the nucleus [45]. Moreover, it exists co-localization of GABA and serotonin at the same neuron[45, 68, 69]. Additionally, other neurotransmitters like glutamate are also present in the MRN [45].

### 4.1 Classic characterization of raphe neurons

Most of the studies that contributed to the characterization of serotonergic neurons have been carried out in the DRN. Early studies detected regularly spontaneously firing neurons in the DRN in anesthetized rats using extracellular recording techniques [33, 35, 66, 67]. Serotonergic neurons of the DRN characteristically show spontaneous activity with a slow (1-5 *Hz*) and regular discharge frequency described as clock-like [66]. This unique firing pattern would serve as a “neuronal signature” for this neurochemical group of brainstem cells [1]. The extracellular action potentials (APs) of these neurons exhibit a prominent positive deflection, followed by a negative or negative/positive deflection. The first positive deflection plus the first negative deflection has a duration greater than 1.4 *ms* [32, 33].

Hajos et al. in 1995 [32] described that during their regular discharge these neurons show spike bursts (from 2 to 4 spikes) [31, 32]. In these bursts, the spikes have a short interval (range: 2.4-11.5 *ms*), and the secondary spikes show a decrease in amplitude. In addition, non-serotonin neurons with varied electrophysiological properties are also present in the DRN. This heterogeneous population of neurons show a less regular pattern of firing, with firing rates in the range of 0.1-30 *Hz* [33–35, 66].

In contrast to the DRN, few studies describe the electrophysiological characteristics of the MRN. These studies were carried out using *in vitro* [45, 70, 71] or anesthetized animals *in vivo* [32, 36–38]. The studies carried out by Kocsis et al.(2006) [37] and Viana di Prisco (2022) [38] showed a great diversity in the MRN neurons: a group of serotonergic neurons which characteristics similar to the “Clock-like” neurons of the DRN, a fast-firing serotonergic group of neurons rhythmic with theta, and a more heterogeneous non-serotonergic group of neurons. Therefore, the electrophysiological criteria used in previous studies to identify “putative” serotonergic neurons would be appropriate only as a rough preliminary classification of MRN neurons emphasizing the need for new metrics for the classification of neuronal activity.

### 4.2 Non-linear metrics in neuronal characterization

Nonlinear metrics had been used to compare neuronal dynamics in different neurological pathologies [72], finding that their values change in different states [73, 74] and reporting that neuronal entropy depends in the level of alertness in humans and in animals. It was also recently used to compare informational complexity in spike trains across species [75]. Another study [76] compared FR with the Shannon Information Transmission Rate in the responses of lateral geniculate nucleus neurons of the cat to spatially homogeneous spots of various sizes with temporally random luminance. They found that the behavior of these two rates can differ quantitatively. This suggests that the energy used for spiking does not translate directly into the information to be transmitted. They also compared FR with Information Rates for two type of cells in the lateral geniculate nucleus: X-ON (neurons excited by light onset) and X-OFF cells (neurons excited by light offset). They found that, for X-ON cells the FR and Information Rate often behave in a completely different way, while for X-OFF cells these rates are much more highly correlated. This results suggest that for X-ON cells a more efficient “temporal code” is employed, while for X-OFF cells a straightforward “rate code” is used.

A recent work by Estarellas et al. (2020) [77], used nonlinear metrics to investigate the encoding and information transmission in time series of sensory neurons. They found that depending on the frequency, specific combinations of neuron/class and coupling-type allow a more effective encoding, or a more effective transmission of the signal. However, our work is the first study proposing non-linear metrics for neuronal characterization of the raphe nuclei using novel informational complexity measures.

We show that compared to the other metrics, the OP Entropy separates better the DRN and MRN groups, pointing to OP Entropy being a strong candidate for neuronal activity categorization.

When performing a neuronal classification, we need to capture different information about the studied phenomenon and superimpose little with the other metrics – a measure that captures the same information than another measure is redundant and not very informative. When we look at the classic measures that are normally used to analyze extracellular neuronal activity, the APD gives a very different information compared to the FR or CV. However, FR and CV are related to each other because the CV is found from the FR and is not independent, which can explain why those two measures are linearly related (Fig. 4A), while the combination of APD with FR or CV are not (Fig. 4B and C).

When we look at the relationship between the classical metrics and the non-linear metrics, a significant linear relationship would be unexpected. For example, the APD (Fig. 4D, E and F) is not significantly related to any of the non-linear metrics. However, when we look at the CV, a strong negative linear relationship is present with the Bins Entropy (Fig. 4H). Those variables could be related by the standard deviation (SD) if the data by which the Bins Entropy is calculated follows a Gaussian distribution. On the other hand, for a given FR, if the SD is broad, the CV will be low and tend to a uniform distribution, and the entropy of a uniform distribution is maximum.

OP Entropy shows a weak or no linear relationship with other metrics studied (Table 1 and Fig. 44), indicating that the information captured by this metric superimpose little with the other metrics. On the other hand, PLZC correlates with the FR (Table 1 and Fig. 4K). Additionally, Bins Entropy and PLZC show a strong positive linear relationship despite they are not redundant.

We also show that OP Entropy is insensitive to the rhythmic patterns present in the time series, whereas all the other metrics change depending on whether the neuronal activity is rhythmic or not (Fig. 5). Additionally, none of the non-linear metrics is sensitive to burst patterns.

Regarding the characterization of putative serotonergic neurons using non-linear metrics, we found that these neurons display a high Bins Entropy (Fig. 6B). That could be explained because the putative serotonergic group were selected by a low CV, which as can be seen in Fig. 4H is negatively related with the Bins Entropy.

Finally, it is important to note that all the complexity analysis performed in this work were done using a *D* = 3. We were not able to test whether the differentiation improved if the *D* value was higher (like 4 or 5) because we are limited in the length of the time series. It is probable that a larger *D* may reveal a stronger differentiation in MRN and DRN as more information would be included in the patterns to calculate the non-linear metrics.

We conclude that non-linear metrics, and specially OP Entropy, contribute significantly to enrich the characterization of raphe nuclei neurons and is a promising metric to distinguish sub-populations based in neuronal dynamics.

## 5 Acknowledgements

This study was supported by the “Programa de Desarrollo de Ciencias Básicas, PEDECIBA” and the “Comisión Sectorial de Investigación Científica” (CSIC) I + D-22620220100148 grant from Uruguay.

